# NRPS-gene disruption in the apple-pathogen, *Neonectria ditissima*

**DOI:** 10.1101/2021.04.01.438028

**Authors:** Heriberto Vélëz, Salim Bourras, Larisa Garkava-Gustavsson, Kerstin Dalman

## Abstract

Apple production in Sweden and elsewhere is being threaten by the fungus, *Neonectria ditissima*, which causes a disease known as Fruit Tree Canker. The disease can cause extensive damages and the removal of diseased-wood and heavily infected trees can be laborious and expensive. Currently, there is no way to eradicate the fungus from infected trees and our knowledge of the infection process is limited. Thus, in order to target and modify genes efficiently, the genetic transformation technique developed for *N. ditissima* back in 2003 was modified. We report on the upgraded protocol and show that protoplasts were viable, able to uptake foreign DNA, and able to regenerate back into a mycelial colony, either as targeted gene-disruption mutants or as ectopic mutants expressing GFP.

## Introduction

The apple industry is the largest fruit sector in Sweden, which has seen a steady increase on the number of hectares used for apple growing by as much as 12% during 2012-2017 (Sandberg and Monica n.d.). However, apple production in Sweden and elsewhere is being threaten by the fungus, *Neonectria ditissima*, which causes a disease known as Fruit Tree Canker (FTC)(R.W.S. Weber 2014) as the fungus can also attack other fruit trees such as pear (*Pyrus communis*) and quince (*Cydonia oblonga*). The cool and rainy climate in Sweden can favoured the fungus, which can cause extensive damages as the diseased-wood must be removed and heavily infected trees must be taken out. In fact, the removal of entire orchards has been necessary in the USA (Jones and Aldwinckle 1990), and in Sweden, more than 10% of trees can be lost due to FTC (Äppelriket, pers.comm.). Taking into account the high costs for establishing a new orchard (∼400,000 SEK/ha), the disease is a serious threat to Swedish-apple production.

*N. ditissima* can infect plants through wounds (e.g., leaf and fruit scars, pruning wounds, broken branches) throughout the year and can cause rotting of fruits in storage (Xu and Robinson 2010; R.W.S. Weber 2014). Fungal spores spread aerially and by rain splashing within the tree and to nearby trees. Branches and the trunk can be girdled, leading to death of all proximal parts, and young trees can become infected during propagation (R.W.S. Weber 2014; Roland W. S. Weber and Børve 2021). *Neonectria* spp. *are* also known to attack many different species of forest trees, thus the disease is also known as European canker (M Ghasemkhani et al. 1989). Neonectria-cankers have been recorded on forest trees in Europe and the USA. In Norway for example, a *N. ditissima*-like fungus was found to infect subalpine fir (*Abies lasiocarpa*), White fir (*A. concolor*), Siberian fir *(A. sibirica*), and Norway spruce (*Picea abies*) (Venche Talgø 2012). There are also reports of Neonectria-canker found in Sweden, Denmark, United Kingdom, and Belgium (Schmitz, Charlier, and Chandelier 2017) . Hence, *Neonectria* spp have the propensity to cause damage to European forestry through the extensive scarring and damage that leads to reductions in quality and value of timber (Metzler et al. 2002).

In apple orchards, chemical control against *N. ditissima* are now prohibited in Sweden, which underscores the need to breed for resistant apple cultivars. Disease control measures rely primarily on the careful monitoring and removal of infected plants, which are laborious and expensive (R.W.S. Weber 2014). Currently, there is no way to eradicate the fungus from infected trees and our knowledge of the infection process by the fungus is limited. Recently, the genome of *N. ditissima* has been sequenced and, thus far, three draft genomes have been published revealing a genome size of about 44 Mbp (Deng et al 2015, Gomez-Cortecero et al. 2015). A genetic transformation system using protoplasts of *N. ditissima* was developed in 2003 using a no-longer-available enzyme mixture, Novozymes 234 (Tanguay et al 2003). Hence, we set out to develop a new genetic transformation system that, together with the genome information available, will enable us to elucidate and increase our knowledge of the infection process. Here we report on the refinement of the established transformation system, the generation of mutants to express the green fluorescent protein (GFP) and to delete AK830_g4721T0, which encodes a nonribosomal peptide synthetase (NRPS). This NRPS is similar to one that was shown to be differentially expressed during fungal infection of the fungus, *Valsa mali*, on apple (Ke et al. 2014), and thus selected for deletion. These mutants will be used on apple-fungal pathogenicity assays to begin elucidating the role of NRPS and the secondary metabolites produced by the fungus during plant-pathogen interactions.

## Materials and Methods

### Strains and culture conditions

Cultures of *Neonectria ditissima* isolate E1 (Marjan Ghasemkhani et al. 2016), were maintained in ASAWA media (Amponsah, Walter, and Scheper 2014). Liquid Modified Melin Norkrans media (MMN; Marx, 1969) prepared with sucrose instead of glucose, was used to grow the fungus for genomic DNA extraction or for the growth of mycelial mass for protoplast generation.

### Identification of the NRPS genes

Blast homology searches against the published genome of *N. ditissima* using the comp24091_c0_seq1 encoding a putative NRPS in the fungus *Valsa mali* (Ke et al. 2014), was used to identify the gene encoding a similar NRPS in the published genome of *N. ditissima* (Gómez-Cortecero, Harrison, and Armitage 2015).

### DNA extraction, PCR and Cloning

DNA was extracted using the NucleoSpin Plant II kit (Macherey-Nagel GmbH & Co. KG, Düren, Germany) and following their protocol. All PCR reactions were done using Phusion Hot Start II DNA Polymerase (Thermo Scientific) following the manufacturers’ protocol, with annealing temperatures at 62°C. Primers (Table 1) were purchased from Integrated DNA Technologies (Belgium) and dNTPs were purchased from Thermo Scientific. All PCR products were gel-purified using the GeneJET Gel Extraction Kit (Thermo Scientific). pJET 1.2 (Thermo Scientific) and One Shot® TOP10 Chemically Competent E. coli (Invitrogen) were used for routine cloning. Plasmids were isolated using the GeneJET Plasmid Miniprep Kit (Thermo Scientific).

**Table 1.**
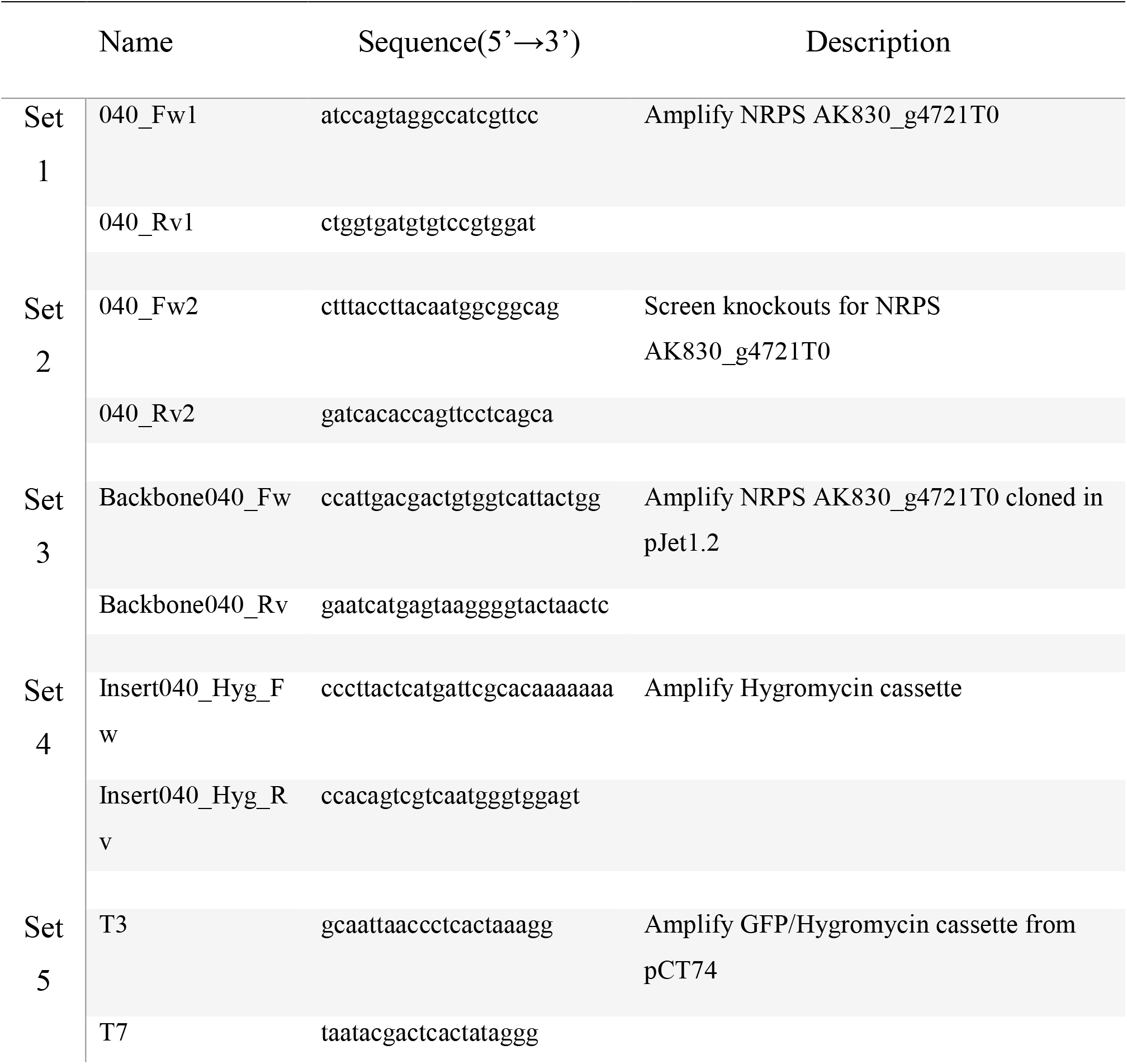
Primers used in this study

### Knock-out Construct generation

Primer Set 1 (Table 1) was used to amplify the coding sequence (3803bp) from the NRPS gene AK830_g4721T0. After gel purification, the PCR product was cloned into pJet1.2 and transformed into One Shot® TOP10 Chemically Competent *E. coli* (Invitrogen). Positives clones were verified by restriction digest. A positive clone from the plasmid prep was used as a template in a PCR reaction with Primer Set 3 (Table 1) to amplify a linear fragment comprising of the backbone of the plasmid flanked with a 5’-fragment (1100bp) and a 3’-fragment (1200bp) from the coding sequence at each end. Primer Set 4 (Table 1) was used to amplify the hygromycin cassette from plasmid pJCA-HygII (Vélëz et al, unpublished). After gel purification, both PCR fragments were used with GeneArt® Seamless Cloning and Assembly Enzyme Mix (Invitrogen) by following the manufacturers’ protocol and at the end of the 30 min incubation step, 5 µL were transformed into One Shot® TOP10 Chemically Competent *E. coli* (Invitrogen). Positives clones were verified by restriction digest.

### PCR products for transformation

A PCR fragment containing the assembled construct (Fig. 1), was amplified from a positive clone using primer Set 1, purified using the E.Z.N.A.® Cycle-Pure Kit (Omega Bio-tek), and used for the transformation of protoplasts to make AK830_g4721T0-knockouts. Similarly, primer Set 5 (Table 1) was used to amplify the coding sequence (3117 bp) for the GFP/Hygromycin cassette from plasmid pCT74 (Lorang et al. 2001) (Fig. 2), purified using the E.Z.N.A.® Cycle-Pure Kit (Omega Bio-tek), and used for the transformation of protoplasts to make GFP-expressing mutants.

**Figure 1.**
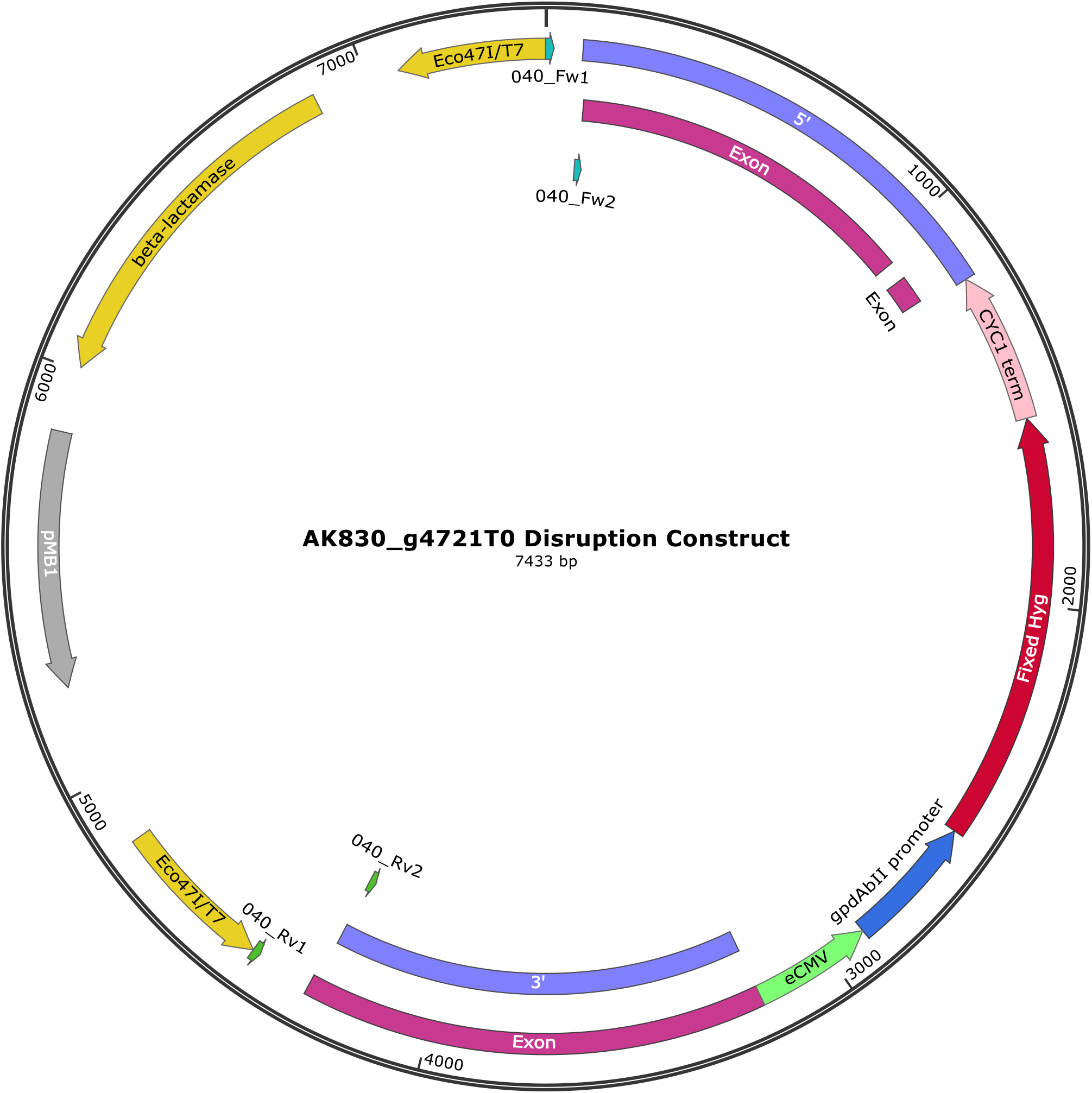
Disruption construct generated by GeneArt (Thermo Scientific). Primers 040_Fw2 and 040_Rv2 (Table 1) were used to amplify 4200 bp comprising the 5’ and 3’ regions flanked by the Hygromycin cassette. Primers 040_Fw2 and 040_Rv2 (Table 1) were used to screen for mutants. Generated using SnapGene® software (from Insightful Science; available at snapgene.com).

**Figure 2.**
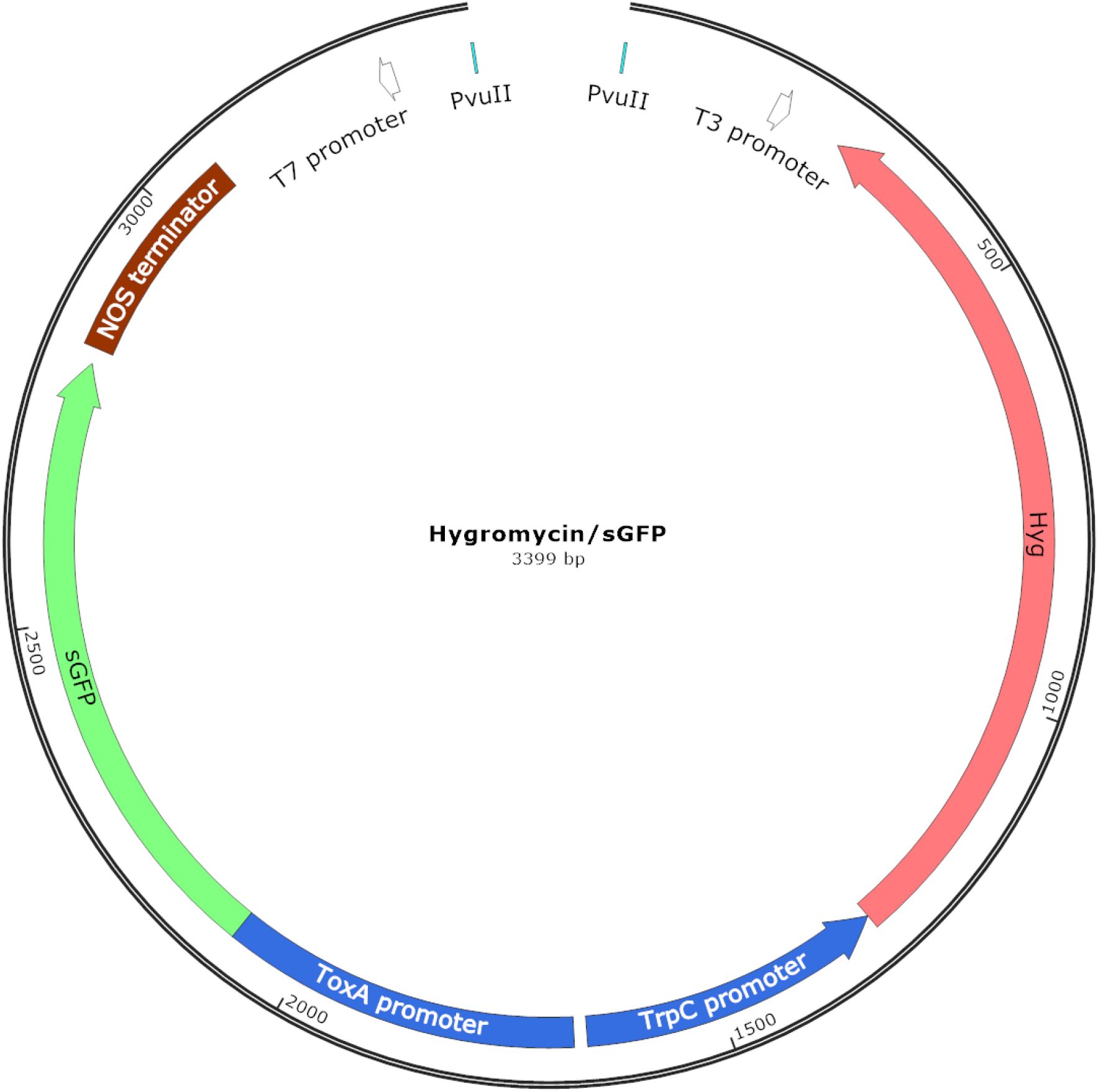
The GFP expression vector, pCT74, developed by Lorang et al., (2001) was used during the transformation. The primers T7 and T3 were used to amplify 3117 bp containing the coding sequence for the GFP expression and hygromycin selection, used during the transformation. Generated using SnapGene® software (from Insightful Science; available at snapgene.com).

### Protoplast production and transformation

To produce protoplasts, the protocol by Tanguay et al., (2003) was followed with several modifications. Spores (10^9^) were transferred to a 250 ml Erlenmeyer flask containing 100 ml of liquid MMN media and incubated at 25°C for four days without agitation. The fungal mycelium was transferred to a 50 ml conical tube and pelleted by centrifugation at 3000 x g at 4°C for 10 min. After decanting the growth media, the mycelium was rinsed with osmoticum buffer (OB; 0.8 M NaCl; 50 mM Maleate Buffer (pH 5.5)) and centrifuged as before, repeating this step once again. Finally, the pellet was re-suspended in four millilitres of filtered-sterilized (0.22 µm filter) lysing buffer (75 mg/mL BSA; 10 mg/mL Yatalase (Takara); 5 mg/mL Lysing Enzymes (L1412 Sigma); 5 mg/mL L-cysteine dissolved in OB), vortexed twice for 30 sec to mix, and incubated for 1.5 hr in a water bath at 30°C with gentle shaking. The protoplasts were separated from cell wall debris by filtration through four layers of sterile Precision Wipes (Kimtech Science, Kimberly-Clark) into a new 50 mL conical tube. The protoplasts were washed twice, with 20 ml of ice cold STC buffer [1 M d-sorbitol, 50 mM Tris–HCl (pH 8.0), 200 mm CaCl2], and pelleted by centrifugation at 2500 x g at 4°C for 8 min each time. The protoplasts were re-suspended in STC and their concentration adjusted accordingly. For the transformation, 100µL of protoplasts (10^8^ in STC) aliquoted in 50 mL conical tubes (Sarsted), were incubated on ice for 30 minutes after the addition of 10 µg of PCR’d-DNA construct in 40 µL of STC and 10 µL of Spermidine (10 mM dissolved in STC and filtered-sterilized). After 30 minutes, 600µL of PTC buffer [40% (w/v) PEG4000, 50 mM Tris-HCl (pH 8.0), 200 mm CaCl2] was added dropwise to the tubes and incubated at room temperature for an additional 30 min. Liquid MYGS (1% maltose, 0.04% glucose, 0.04% yeast extract, and 0.7 M sucrose) was added to the 20 mL mark and placed at 28°C for 24 hrs. The following day, each 50 mL conical tube was subdivided into two 10 mL-aliquots and each aliquot was mixed with 30 ml of ASAWA media (containing 0.7 M sucrose and 1% (w/v) low melting point agarose), 30 µg/ml of Hygromycin B and poured into big plates (150 ⱷ; Sarsted) after gentle mixing. Plates were incubated at 28°C for 10 days in the dark and putative transformants (selected as single colonies) were transferred to ASAWA plates.

## Results

### Identification of the NRPS genes

Two genes encoding proteins similar to comp24091_c0_seq1 from the fungus *V. mali* were found in the genome of *N. ditissima*, AK830_g2733T0 and AK830_g4721T0 (Fig. 3). The proteins shared 46% amino acid sequence similarity (64% positives).

**Figure 3.**
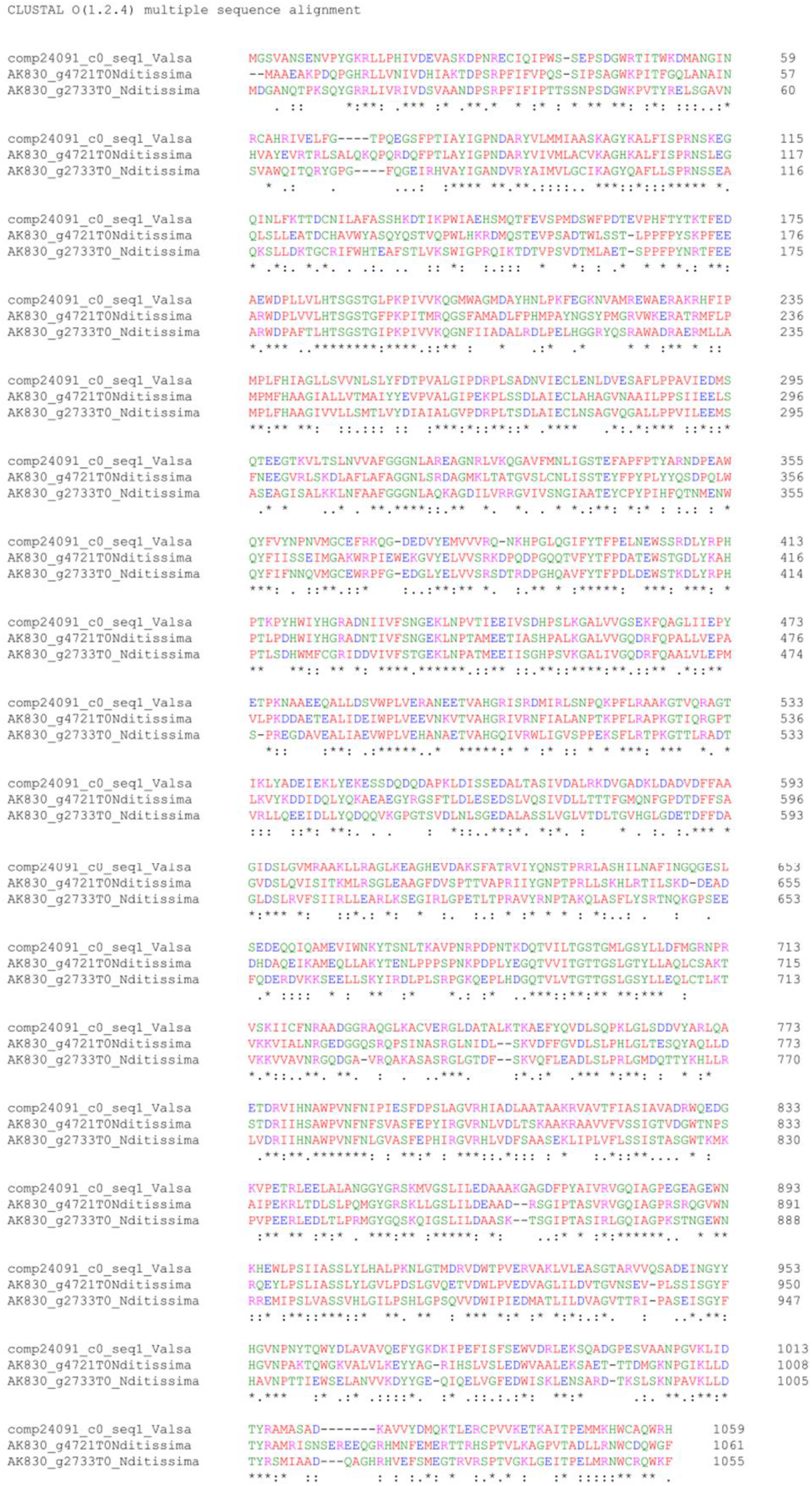
Amino acid alignment of comp24091_c0 from *Valsa mali*, compared to AK830_g2733T0 and AK830_g4721T0 from *Neonectria ditissima*. The alignment was generated using Clustal Omega (www.ebi.ac.uk/Tools). Mutants for AK830_g4721T0 were generated in this study.

### Gene knockouts and GFP mutants

Hygromycin resistant colonies were recovered from three independent experiments, approximately 6 to 10 days after transformation. A summary of the mutants generated is presented in Table 2. Approximately, 10 to 50 colonies from gene knockout experiments were recovered (targeted homologous recombination), while there were fewer colonies generated when using only the GFP construct (ectopic integration). PCR screens confirmed the disruption of gene AK830_g4721T0 in the mutants (Fig. 4). Visualization of GFP-expressing mutants was done using a Leica DM5500B microscope equipped with a DFC360fx camera.

**Table 2.** Summary of mutants generated in this study.

| Gene | Selective marker | Colonies picked | Gene deletion |
| --- | --- | --- | --- |
| AK830_g4721T0 | Hygromycin | 15 | 15 |
| GFP Transformation 1 | Hygromycin | 10 |  |
| GFP Transformation 2 | Hygromycin | 5 |  |

**Figure 4.**
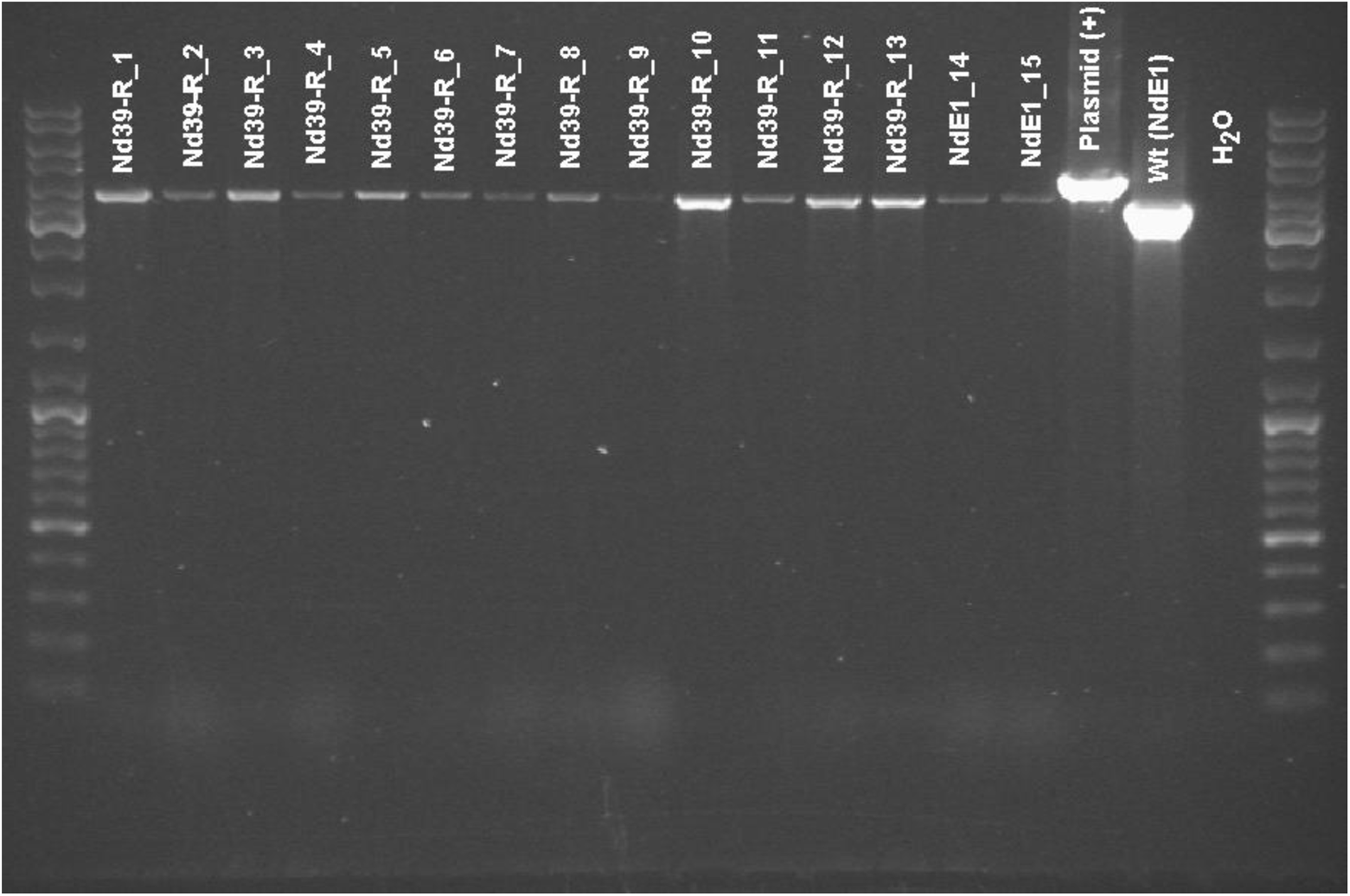
PCR screen of mutants and wild type *Neonectria ditissima*. Primers 040_Fw2 and 040_Rv2 (Table 1) were used to screen for mutants. A size increase from 3548 bp to 4459 bp would be indicative of a gene disruption. Plasmid DNA, Wild Type genomic DNA, and water were included as controls. A 1 Kb ladder (Thermo Fisher) was included in the first and last lane.

## Discussion

In the initial protocol by Tanguay et al., (2003), the enzyme mixture Novozym 234 used in the production of protoplasts is no-longer-available. Thus, a new enzyme mixture called Yatalase (Takara) was prepared and tested with and without Lysing Enzymes (L1412 Sigma). Protoplasts from both tests were viable and able to regenerate. However, protoplasts stored at -70°C and used in later transformations were inviable (data not shown).

To test the ability of the protoplasts to uptake foreign DNA, PCR products for either ectopic integration (to express GFP) or targeted gene-disruption (to knockout genes), were used to treat the protoplasts. Selection of transformed and non-transformed protoplast at 25, 30, and 35 µg/ml of hygromycin, showed that 25 µg/ml was enough to kill wild type and select for mutants (data not shown). However, individual colonies were always picked from the 35µg/ml hygromycin-containing plates. Similarly to Tanguay et al., (2003), transformation varied between experiments. Ectopic transformation with the GFP construct, generated fewer colonies than when targeted gene-disruption via homologous recombination was the intention (Table 2). Nevertheless, we were able to generate 15 mutants showing that gene AK830_g4721T0, had been deleted (Fig. 4), while GFP-expressing mutants under the microscope were clearly “glowing” (Fig. 5). AK830_g4721T0, encodes a NRPS, similar to the one shown to be differentially expressed during the fungal infection of the fungus, *Valsa mali*, on apple (Ke et al. 2014). Thus, these mutants will be used in pathogenicity assays with apple cultivars to further expand our knowledge of the infection process during host-pathogen interactions.

**Figure 5.**
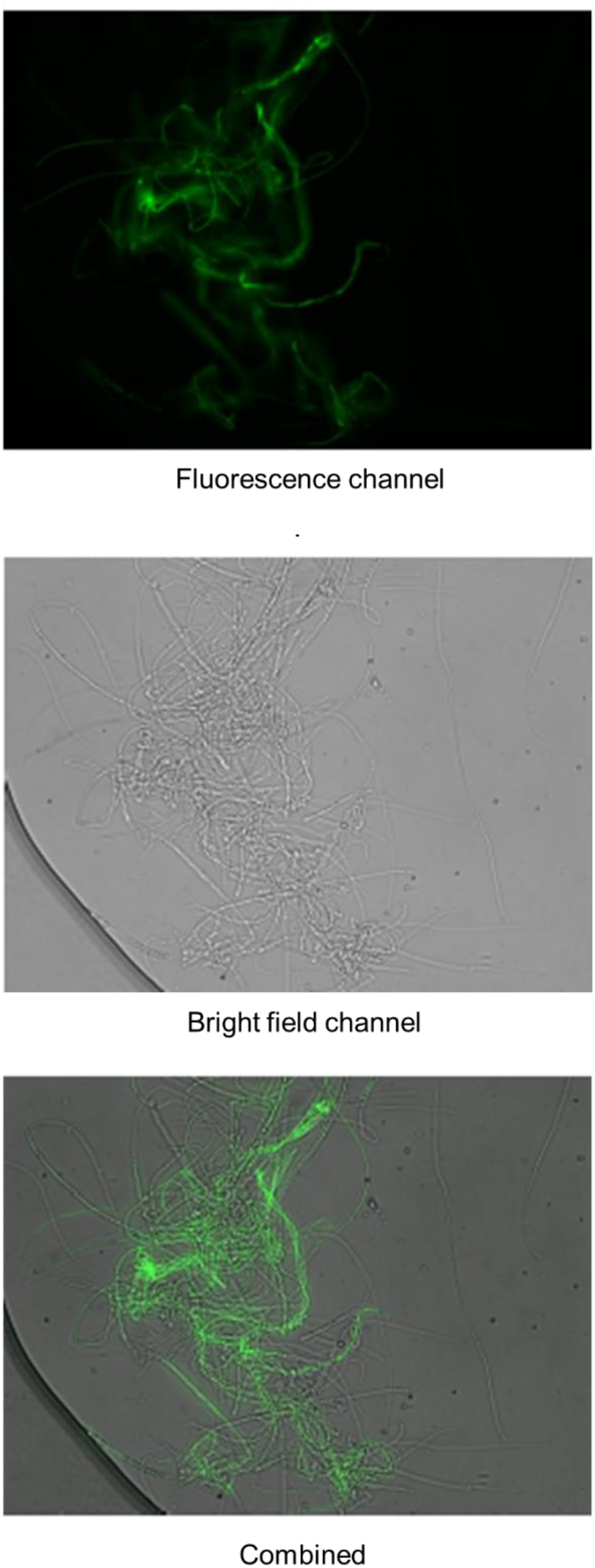
*Neonectria ditissima*-mutant expressing the green fluorescent protein (GFP). Pictures were taken using a Leica DM5500B microscope.

## Conclusion

We have upgraded the protocol established by Tanguay et al., (2003) to make protoplast of the pathogenic fungus, *Neonectria ditissima*, using currently available protoplasting enzymes. The protoplasts were viable, able to uptake foreign DNA, and able to regenerate back into a mycelial colony, either as targeted gene-disruption or as ectopic mutants expressing GFP.

